# Modulation of Retinal Inflammation Delays Degeneration in a Mouse Model of Geographic Atrophy

**DOI:** 10.1101/2023.02.08.527757

**Authors:** Raela B Ridley, Brianna M Bowman, Jieun Lee, Erin Walsh, Michael T Massengill, Alfred S Lewin, Cristhian J Ildefonso

**Author notes:** **Corresponding Author:** Cristhian J Ildefonso, Ph.D., Research Assistant Professor Department of Ophthalmology, University of Florida College of Medicine, PO Box 100284, Gainesville FL 32610. These authors contributed equally to this work (Raela B Ridley, Brianna M Bowman).

## Abstract

The advanced form of AMD, geographic atrophy, is associated with increased RPE oxidative stress and chronic inflammation. Here we evaluated the effects of delivering an anti-inflammatory viral gene by an AAV-vector in a mouse model of geographic atrophy. We measured changes in retinal function, structure, and morphology over nine months with electroretinography, optical coherence tomography, and fundoscopy, respectively. In addition, we used retinal tissue to quantify changes in markers of inflammation by multiplex ELISA, RT-qPCR, and immunofluorescence staining. Our AAV significantly delayed the loss of retinal function and structure and decreased retinal inflammation compared to the control AAV treatment. Our results suggest that modulating retinal inflammation could significantly slow the progression of geographic atrophy.

## INTRODUCTION

Age-related macular degeneration (AMD) is the principal cause of vision loss in developed countries among individuals 60 years old and older^1^. Affected individuals accumulate deposits that can progress from “soft” to “hard” drusen. The presence of complement proteins and complement-associated proteins in drusen deposits and genome-wide association studies (GWAS) point to an inflammatory process^2^. GWAS studies have identified variants of complement factor H and the ARMS2/HTRA1 locus, which is strongly associated with the development of AMD^3^. More recently, the expression of complement factor-related genes has been associated with AMD, further strengthening the role of inflammation in the disease^4^.

Viruses have evolved to evade their host’s inflammatory response. The myxoma virus is a poxvirus from the Leporipoxvirus genus. In this virus, the deletion of the M013 gene permits the detection of the virus by the host immune cells and triggers an inflammatory response^5^. The M013 protein contains a pyrin domain that inhibits the NLRP3-inflammasome signaling pathway. Its C-terminal domain also inhibits the nuclear translocation of p65 (an NF-κB subunit)^6,7^, therefore blocking the expression of potent pro-inflammatory genes such as TNF and interleukin-1 beta (IL-1β).

Our group has developed an adeno-associated viral (AAV) vector that delivers a secretable and cell-penetrating form of the myxoma M013 gene^8^. This vector significantly decreased the number of infiltrative cells within the vitreous humor of an animal model of experimental autoimmune uveitis^9^. We also demonstrated that this vector could considerably reduce the concentration of IL-1β in the vitreous humor of these animals.

Given the activation of inflammatory pathways (such as the inflammasome) in AMD^10–15^, we tested the effects of a single intravitreal injection of our AAV delivering the secretable and cell-penetrating form of M013 in the RPE-specific *Sod2* knock-out mouse model of geographic atrophy (an advanced form of AMD)^16–18^. This model renders the RPE more susceptible to oxidative damage and recapitulates some cardinal features of the human disease, including increased lipofuscin autofluorescence, disruption of Bruch’s membrane, accumulation of sub-RPE deposits, and deposition of complement proteins. These changes can be documented in living mice by declines in the electroretinography (ERG) responses, fundus abnormalities, and alterations to photoreceptor and RPE reflectance measured by spectral-domain optical coherence tomography SD-OCT. The current study found that treating these mice with AAV expressing the secreted and cell-penetrating M013 gene reduced inflammation and delayed retinal degeneration in these mice.

## MATERIALS AND METHODS

### Chemicals and Antibodies

Antibodies used and their dilutions are given in **Table 1**. Antibodies were stored at −4⁰C. The sequences of primers provided in Supplementary Table 1. All primers were synthesized by Eurofins Genomics, resuspended in RNAse and DNAse-free water, and stored at −20⁰C.

### Animals

The University of Florida IACUC approved the experiments described in this work. All procedures adhered to the ARVO guidelines for the use of animals in biomedical research. Mice were reared in a 12 hrs light and 12 hrs darkness cycle. C57BL6J mice (8-19 weeks of age) were use in experiments involving the endotoxin-induced uveitis model as described below. For studies focused on geographic atrophy, we used the RPE-specific Sod2 conditional KO mouse model previously described^16,17,19,20^. We fed mice with doxycycline-containing chow (200 mg doxycycline/diet kg) for two weeks. We then switched to regular chow for the rest of the experiment. Before each procedure, mice were anesthetized with an intraperitoneal injection of ketamine/xylazine mixture (100 mg/kg ketamine, 4 mg/kg xylazine). After the procedure, mice received an intraperitoneal injection of atipamezole (1 mg/kg) to reverse the effects of the anesthesia. Animals were kept on a heated plate until fully ambulatory before returning to racks.

### Vector Design

We created the sGFP-TatM013v5 vector by fusing the sequence of the cell penetrating peptide of HIV Tat and the M013 gene of myxoma virus containing a V5 epitope tag. We cloned this fusion downstream of the IgG kappa light chain leader sequence linked in-frame to the coding sequence of green fluorescent protein (GFP). This gene cassette was then cloned in the pTR-smCBA AAV plasmid (ref) (Fig. 1A). The pTR-smCBA-Igκ-GFP-TatM013v5 plasmid was packaged, purified on Iodixanol gradients, and tittered by the Ophthalmology Vector Core at the University of Florida according to published methods^21,22^.

**Figure 1.**
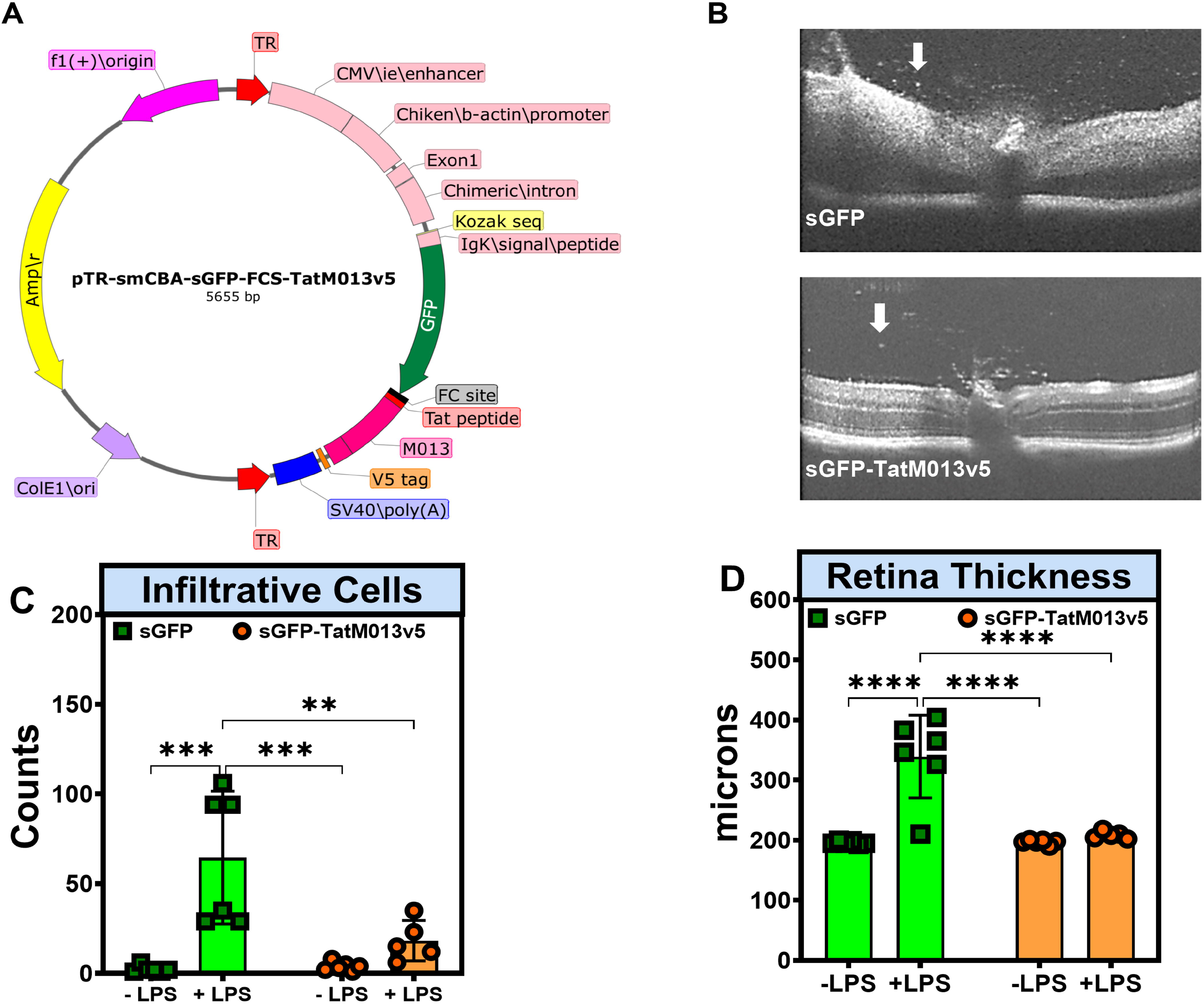
TatM013v5 decreases ocular inflammation in the EIU mouse model. **(A)** Genetic map of the pTR-smCBA-sGFP-FCS-TatM013v5 vector. The control vector has the same sequence except with the TatM013v5 insert. **(B)** Representative SD-OCT B-scans of mice treated with AAV that delivered either sGFP or sGFP-TatM013v5 twenty-four hours after endotoxin (LPS) injection. **(C)** Quantification of infiltrative cells in the vitreous humor was done by two-blinded individuals using Image J software. LPS injection only significantly increased infiltrative cells in sGFP treated eyes. **(D)** Total retinal thickness measurements were done before and after the LPS injection. LPS injection caused a significant increase in retina thickness only in sGFP treated eyes. Values represent average ± standard deviation (n=5-6 mice). ** = adjusted p-value ≤ 0.01, *** = adj. p-val ≤ 0.001, **** = adj. p-val ≤ 0.0001

### Intravitreal injection of AAV

The eyes of mice were dilated using ophthalmic solutions of atropine (1%) and phenylephrine (2.5%). Next, mice were anesthetized with a ketamine/xylazine mixture (100 mg/kg ketamine, 4 mg/kg xylazine). A drop of proparacaine hydrochloride ophthalmic solution (0.5%) was added to the eye as a topical anesthetic. An incision was made in the limbus region using a 27 G needle, followed by inserting a Hamilton syringe with a 33 G needle. Once inside the vitreous, as observed through a Leica stereoscope, 1 µL of the appropriate AAV was injected. Afterward, the needle was retracted, and the mice received an intraperitoneal injection of Antisedan® (1 mg/kg) and topical antibiotics at the injection site. Mice were placed on a warm pad until ambulatory before being returned to their housing rack.

### Endotoxin-induced uveitis (EIU) model

We used the EIU mouse model to validate the anti-inflammatory properties of the V5-tagged M013 gene as done previously^8^. Briefly, C57BL/6J mice were injected with the corresponding AAV vector. In addition, we injected mice intravitreally with 125 ng of lipopolysaccharide (LPS) per eye four weeks after the vector injection.

### Spectral Domain Optical Coherence Tomography (SD-OCT)

Mice eyes were dilated with topical atropine and epinephrine solutions and anesthetized with a ketamine-xylazine mixture (100 mg/kg ketamine: 4 mg/kg xylazine) given intraperitoneally (i.p.). Anesthetized mice were placed on a platform and restrained in place. Using a Bioptigen spectral domain optical coherence tomography equipment, a total of 250 B-scans centered on the optic nerve head were obtained. Afterward, mice received an i.p. injection of atipamezole (0.25 mg/kg) and were kept on a heated pad until ambulatory. Scans were averaged into 25 B-scans subjected to auto-segmentation using the Diver Software from Bioptigen.

### Fundoscopy

‘The eyes of mice were dilated with drops of phenylephrine and atropine ophthalmic solutions. Mice were then anesthetized with a mixture of ketamine/xylazine, as described earlier. Next, mice were placed on a platform, and their eyes were covered with a drop of Genteal to avoid dehydration. A micron III fundus camera was focused on the retina centered about the optic nerve. Images were taken under bright fields and fluorescent filters. The same exposure time and light intensity were used for all the images.

### Electroretinography (ERG)

Mice were dark-adapted overnight. The following day, the ‘eyes of mice were dilated with eyes drops of atropine and epinephrine, as described earlier. Next, mice were placed on a platform and electrodes on their mouth, tail, and corneas. Corneal electrodes were adjusted until similar impedance values (<10 µV) were obtained for both eyes. In comparison, mouth and tail impedance were <16 µV. Mice were then subjected to three flashes of 20 cds/m2 separated by 2 minutes intervals using a Ganze field dome. After the last recording, values were averaged, and the a-, b-, and c-wave amplitude were determined. Finally, mice received an i.p. injection of atipamezole as described earlier.

### Retina RNA Extraction

Mice were humanely euthanized, and their retinas were extracted and placed in 0.4 mL of TriZol reagent. An equal volume of 100% ethanol was added to the samples and mixed. We used the Direct Zol miniprep kit from Zymo Research (Irvine CA) to isolate the RNA per the manufacturer’s protocol. RNA concentration was quantified by Qubit 2.0 as per the manufacturer’s protocol. Samples were stored at −80°C.

### Real-time quantitative PCR (RT-qPCR)

As per the manufacturer’s protocol, a cDNA library was generated using 500 ng of total RNA using the iScript cDNA synthesis kit from Bio-Rad. The concentration of the cDNA was then quantified using the Qubit 2.0 assay for oligos per the manufacturer’s protocol. Samples were then diluted to 2 ng/µL. qPCR reactions were set using 3 µL of each cDNA sample (6 ng), 5 µL of Sosofast Eva Green (Bio-Rad) 2X master mix, and 2 µL of forward and reverse pre-mixed primers (10µM each). Reactions were duplicated for each sample and gene using a hard shell 96-well PCR plate. Primer sequences are given in **Supplementary Table 1**. PCR conditions were as follows: *95°C for 30 seconds, 98°C for 10 seconds, 60°C for 30 seconds, repeat steps 2 and 3 another 39 times, and melt curve from 65-95°C with increments of 0*.*5°C every 5 seconds*. The actin gene primer set was used as the constitutive gene to standardize C_t_ values of each sample. Fold changes were determined using the ΔΔC_t_ method^23,24^. Values were plotted as the Log2 of the fold change.

### Immunofluorescence

Mice were euthanized using CO_2,_ followed by cervical dislocation. Eyes were enucleated, placed in 4% paraformaldehyde in PBS 1X, and kept on ice for 20 minutes. Eyes were then incubated in a 10% sucrose in PBS 1X solution for 1 hour, followed by another hour of incubation in a 20% sucrose in phosphate buffered saline (PBS) solution. Samples were then transferred to a 30% sucrose in PBS and kept at 4⁰C overnight. Eyes were rinsed once in Optimum Cutting Temperature (OCT) medium and then placed in a plastic mold containing the OCT medium. Eyes were then snap-frozen using liquid nitrogen. Fourteen-micron thick sections were collected using a Leica cryostat and fixed onto a glass slide using cold acetone. Slides were stained using a Shandon sequenza slide rack and permeabilized with a 1% Triton-X100 in PBS supplemented with 5% normal horse or goat serum, depending on the antibodies used. Primary and secondary antibodies were used at the dilutions indicated in **Table 1**. DAPI was used at a 1:1,000 dilution as a nuclear counterstain. Finally, slides were imaged using a wide-field Leica fluorescence microscope and the LAS-X software from the same company.

**Table 1.** Antibodies.

| Antibody | Host | Dilution | Company | Catalogue No. |
| --- | --- | --- | --- | --- |
| anti-V5 | mouse | 1 in 5,000 | Invitrogen | R960-25 |
| anti-Tubulin | rabbit | 1 in 1,000 | Novus<br>Biologicals | NB600-936 |
| anti-Rabbit-CW800 | Donkey | 1 in 5,000 | Li-COR | 926-32211 |
| anti-Mouse-CW680 | Donkey | 1 in 5,000 | Li-COR | 926-68070 |

**TABLE 2.**
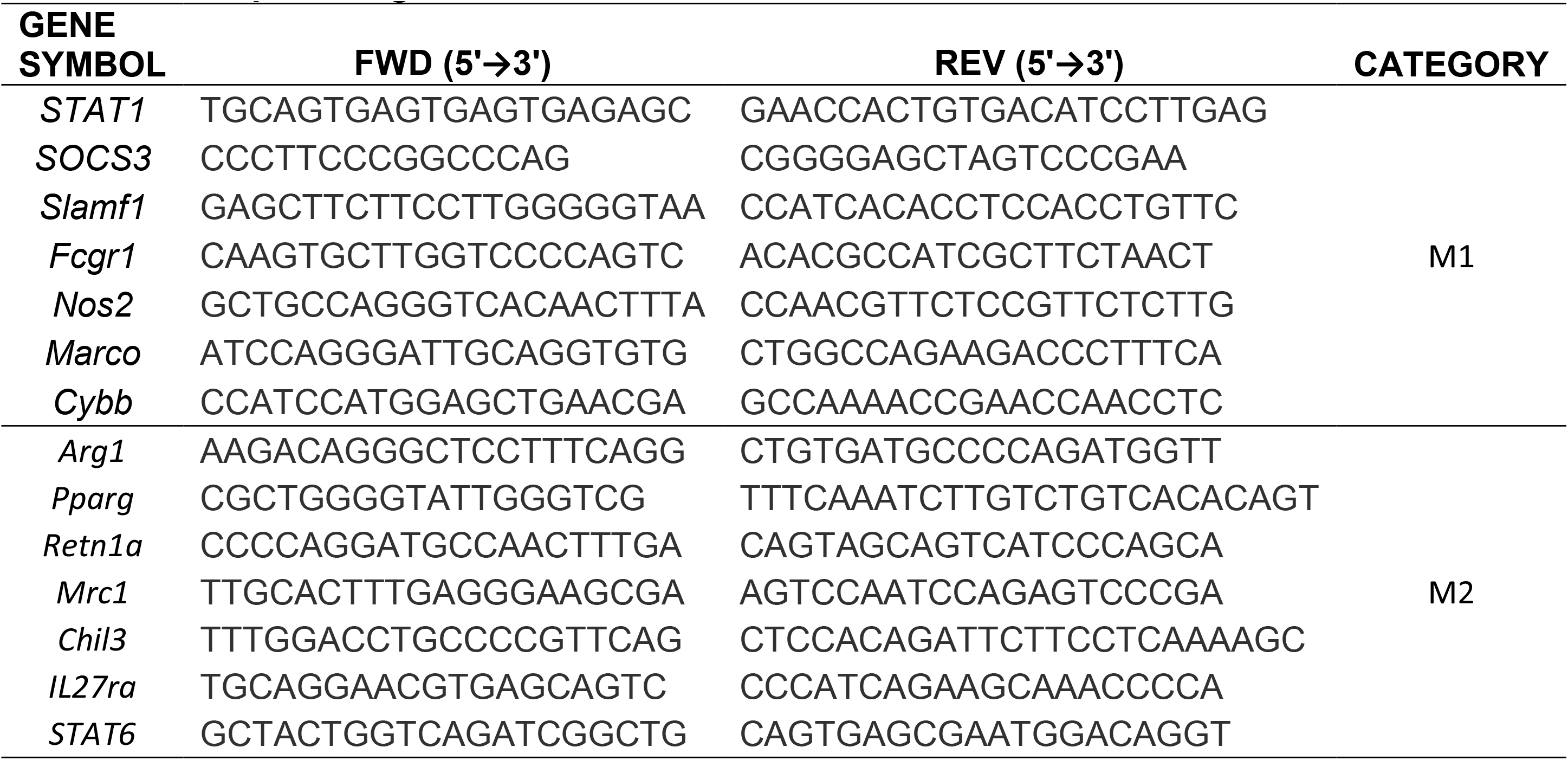
RT-qPCR Oligonucleotides.

### Retinal Protein Extraction

Mice were humanely euthanized by CO_2_ inhalation, followed by cervical dislocation to ensure their death. A horizontal incision was made through the cornea using a number 10 scalpel. The aqueous humor was blotted onto a tissue. The cornea was removed using micro scissors, and the lens, vitreous, and retina were squeezed by applying pressure on the back of the globe with a pair of forceps. The lens and vitreous were separated from the retina, which was then placed on 100 µL of NP-40 lysis buffer (1% NP-40, 50 mM Tris-HCl (pH 8.0), 150 mM NaCl) supplemented with Halt’s protease inhibitors cocktail (Sigma-Aldrich). Retinas were then homogenized with 30 firm strokes of a motorized pestle. The samples were centrifuged at 10,000 x g for 5 minutes at 4°C to remove non-solubilized material. Retina homogenates were collected and transferred into a new and sterile 1.5 mL tube. Finally, protein concentration was determined using the Dc Assay (Bio-Rad) per the manufacturer’s protocol. Samples were then diluted to 1 µg/µL using NP-40 lysis buffer.

### Protein concentration assay

Protein concentration was determined using the Bradford Assay from Bio-Rad. Briefly, BSA concentration was determined with a nano spectrophotometer. Standard was diluted into 1.5, 1.2, 0.9, 0.6, and 0.3 μg/μL using NP-40 lysis buffer. Reagent A from the kit was mixed with reagent S as indicated by the company protocol to make reagent A’. A total of 25 µL of A’ was added to each well, followed by adding 5 µL of the sample. Next, 200 uL of Reagent B was added to each well using a multichannel pipette, followed by a 15-minute incubation at room temperature with constant shaking. Absorbance at 750nm was determined using a plate reader. The absorbance of blank wells was subtracted from all standards and samples. Samples concentrations were extrapolated from a linear regression generated using the BSA standards.

### Multiplex ELISA

A total of twenty-five micrograms of protein lysate were used to set up a multiplex ELISA plate. We used the MILLIPLEX Map Mouse cytokine/chemokine Magnetic Bead Panel-Premixed Plex-Immunology Multiplex Assay (MCYTMAG-70K-PX32) per the manufacturer’s instruction. Briefly, the Mouse Cytokine/Chemokine Magnetic Bead Panel plate was prepared by washing it with 200 µL of Wash Buffer and shaking it vigorously for 10 minutes. The Wash Buffer was decanted, and the residual buffer was removed by lightly tapping the plate on an absorbent towel. 25 µL of the Standards and Quality Controls were duplicated to the appropriate wells. An assay buffer was used as the blank standard. 25 µL of Assay Buffer was added to all wells corresponding to the samples, standards, and quality controls. 25 µL of each sample was added in duplicate to the appropriate wells. The Antibody-Immobilized Pre-mixed Beads were prepared by sonicating the bottle for 30 seconds and then by vortex mixing the bottle for 1 minute. After vortexing, 25 µL of the pre-mixed beads were added to each well. The beads were mixed intermittently by vortexing after every column. The plate was sealed with a foil plate sealer and incubated at 4℃ overnight with agitation on a plate shaker.

The next day, the contents of the wells were removed by placing the plate on a Magnetic 96-well plate Separator (Millipore, Burlington, MA, US) for 1 minute to allow the magnetic particles to remain bound. 200 µL of Wash Buffer was added to the plate, shaken vigorously for 30 seconds, placed on the magnetic plate for 1 minute, and then decanted. This step was repeated twice. After the detection antibodies were allowed to warm to room temperature, 25 µL was added to each well containing the Standards, the Quality Controls, and the samples. The plate was sealed with a foil plate sealer and incubated at room temperature with agitation on a plate shaker for 1 hour. A total of 25 µL of Streptavidin-Phycoerythrin was added to each well containing 25 µL of the Detection antibody. The plate was sealed with a foil plate sealer and incubated at room temperature with agitation on a plate shaker for 30 minutes. The plate was placed on the magnetic plate for 1 minute to allow the magnetic particles to remain bound and then decanted. A total of 200 µL of Wash Buffer was added to the plate, shaken vigorously for 30 seconds, placed on the magnetic plate for 1 minute, and then decanted. This step was repeated twice. Next, 150 µL of Wash Buffer was added to all the wells containing the Standards, the Quality Controls, and the samples. The plate was run on MAGPIX with Xponent Software (Millipore, Burlington, MA, US). A total of 150 beads per region were acquired per each well.

### Western blot

Protein lysates were diluted to 1 µg/µL in NP-40 lysis buffer and 5X Laemmli Buffer^25^ containing 100 mM DTT. Samples were boiled for 5 minutes to denature proteins. A total of 25 µg of protein were loaded into a 4-12% SDS-PAGE gel and ran at 200 volts for 22 minutes. Proteins were transferred onto a nitrocellulose membrane with the iBind 2. The membrane was then incubated with anti-V5, anti-beta tubulin, donkey anti-mouse CW680, and donkey anti-rabbit CW800 using the iBind Flex system from Invitrogen as per the manufacturer’s protocol. Finally, the membrane was scanned with an Odyssey CLx scanner.

### Statistical Analysis

Values are reported as average ± standard deviation. Data were analyzed using one-way, or two-way ANOVA followed by a post-hoc Holm-Sidak test using GraphPad Prism 9 (La Jolla, CA). Statistical significance was considered when either the p-value or adjusted p-value ≤ 0.05.

### Data availability

All the data will be available through the Harvard Dataverse website (https://support.dataverse.harvard.edu)

## RESULTS

### Tagged M013 retains its anti-inflammatory properties

We have previously demonstrated that gene delivery of the myxoma virus M013 gene protects the retina in the endotoxin-induced uveitis (EIU) and autoimmune uveitis mouse models ^8,9^. However, there is currently no antibody to detect the expression of M013. Thus, we added a V5 epitope tag (GKPIPNPLLGLDST) to the C-terminus of the M013 cDNA **(Fig1 A)**. To validate that adding a V5 tag did not affect the function of M013, we tested the sGFP-TatM013v5 AAV vector in the EIU mouse model, which is generated by injection of *E. coli* lipopolysaccharide (LPS) into the vitreous 24 hours before analysis by SD-OCT and sacrifice of the mice. All mice were injected intravitreally with AAV delivering either sGFP or sGFP-TatM013v5 one month before EIU induction with LPS. We quantified the recruitment of inflammatory cells to the vitreous humor and the retina thickness using SD-OCT **(Fig 1B)**.

When the number of infiltrative cells was quantified before and after the intravitreal injection of endotoxin (125 ng LPS), we observed a 28-fold increase in these cells in sGFP-treated retinas. In contrast, retinas treated with the TatM013v5 had a 4.7-fold increase in the number of infiltrative cells. Furthermore, when we compared retinas treated with LPS only, treatment with TatM013v5 significantly decreased the number of infiltrative cells **(Fig 1C)**.

Total retinal thickness was also measured using SD-OCT images. Retinas treated with sGFP showed an average 73% increase in thickness when injected with LPS. In contrast, TatM013v5-treated retinas had only a 5.5% increase in total thickness when treated with the same dose of LPS **(Fig 1D)**. Together, our results demonstrate that our M013 cDNA with a V5 tag retained robust anti-inflammatory properties.

### Intravitreal delivery of anti-inflammatory AAV delays retinal loss-of-function in a GA mouse model

There are multiple mouse models of geographic atrophy; however, none recapitulate all the clinical features of the human disease. To test our gene therapy approach, we used the RPE-specific *Sod2* KO (RPE-Sod2 cKO) mouse model developed by Mao et al.^16^. The RPE-Sod2 cKO mouse is a genetic model in which the Sod2 gene is deleted exclusively within the RPE cells, thus leading to increased oxidative damage and a slow structural and functional decline of the retina^19^. We have previously demonstrated that the RPE-Sod2 cKO mouse model develops slow retinal degeneration and chronic retinal inflammation^26^. We injected two-month-old mice of the correct genotype intravitreally with 10^10^ vector genome copies of AAV2quad(Y-F)+T495V vector delivering either secreted-GFP (sGFP) or sGFP-TatM013v5 under the control of the constitutively active small CBA promoter. We evaluated mice a month after vector injection using fluorescence fundoscopy to validate transgene expression. Mice showed diffused GFP expression in both eyes, thus demonstrating that our AAV vectors express the secreted GFP encoded in their genetic cargo **(Fig 2A-D)**. Furthermore, we detected the secreted and cleaved TatM013v5 in retina protein lysates as a 15 KDa band **(Fig 2E)**.

**Figure 2.**
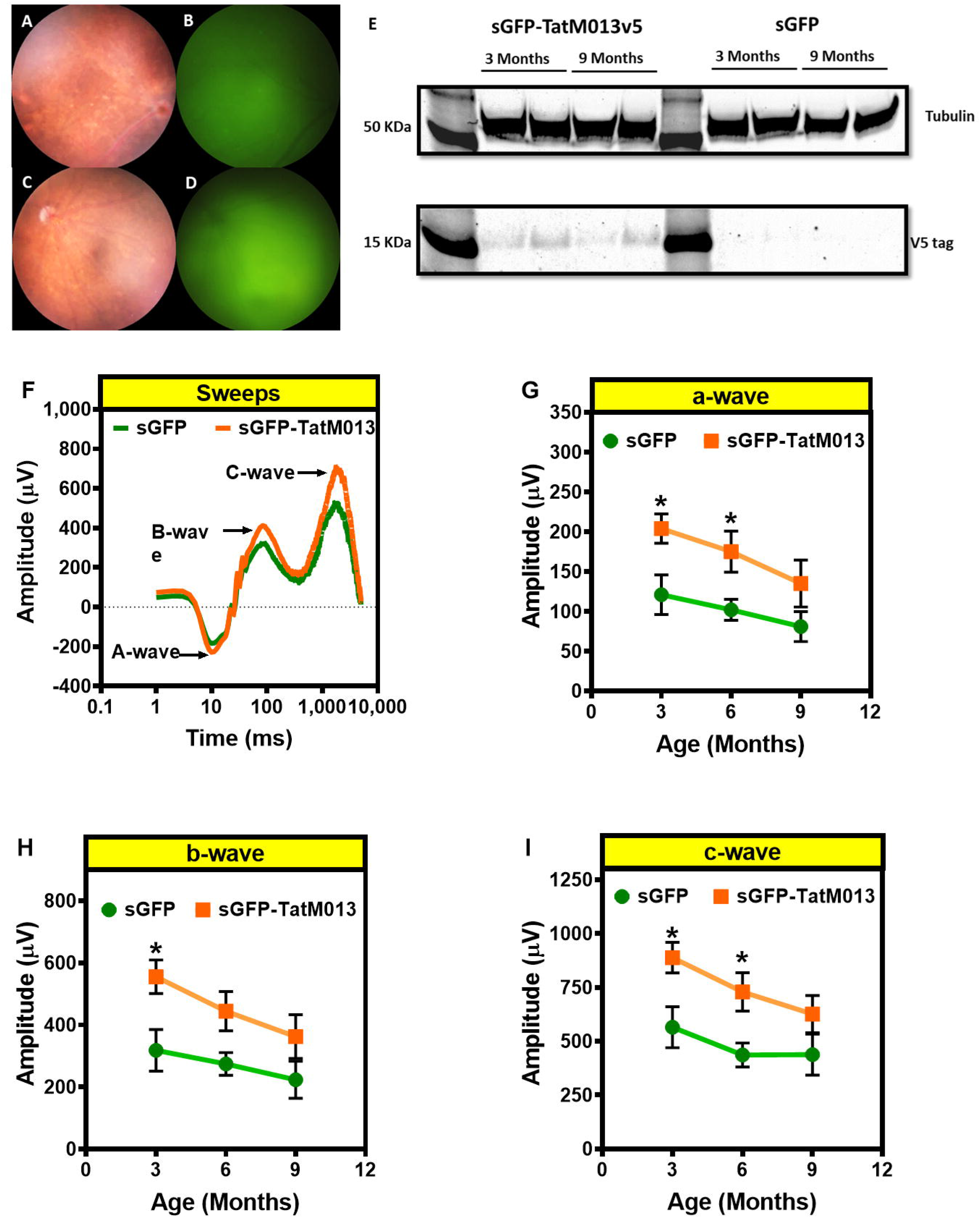
Retinal delivery of TatM013 in a mouse model protects the retina function of the RPE-specific *Sod2* KO mouse model. **(A-D)** Two-month-old mice of the correct genotype were injected intravitreally with 10^10^ vgc of AAV2quad(Y-F)+T491V, delivering either sGFP or sGFP-TatM013. One month later, fluorescent fundoscopy using a Micron III camera confirmed transgene expression. Representative images demonstrate the presence of GFP fluorescence in the retina of either sGFP or sGFP-TatM013-treated eyes. **(E)** Western blot demonstrating the retinal expression of TatM013v5 in sGFP-TatM013v5 treated mice. Mice were evaluated with ERG at 3-, 6-, and 9-months of age. **(F)** Representative ERG recording at 6 months of age. The average amplitudes of their **(G)** a-wave, **(H)** b-wave, and **(I)** c-wave at each time point are graphed as a function of time in months. Eyes treated with the AAV vector delivering sGFP-TatM013 had significantly higher amplitudes when compared to sGFP-treated eyes. Values are reported as average ± standard deviation. (n= 5 mice per group). * = adj. p-val ≤ 0.05

To determine whether the TatM013 vector protected the retina function of our mouse model, we used electroretinography (ERG) to evaluate the sensitivity of the retina to light **(Fig 2F)**. At three months of age, one month after treatment, the amplitudes associated with the activity of the photoreceptors (a-wave), second-order neurons (b-wave), and even the RPE (c-wave) were significantly higher in eyes treated with the TatM013 expressing vector when compared with those treated with sGFP expressing vector. In addition, animals treated with the TatM013v5 vector had a higher a-, b-, and c-wave amplitude at three and six months after injection **(Fig 2G-I)**. Moreover, the average amplitude for all these waves at nine months in TatM013v5-treated animals was similar to the amplitudes for sGFP-treated animals at three months. These observations suggest that although TatM013v5 expression does not stop retinal loss of function, it significantly delays retinal degeneration.

### TatM013v5 treatment preserves photoreceptors’ inner and outer segments

The effects of the retinal expression of TatM013v5 in RPE-Sod2 cKO mice on retinal structure were studied using spectral-domain optical coherence tomography (SD-OCT). We evaluated mice at 3, 6, and 9 months of age to determine if the functional effects observed by ERG correlated with structural effects on their retina. Our results indicate that the retina nerve fiber layer (RNFL) was significantly thicker in the TatM013-treated mice compared to sGFP treated mice at three months of age **(Fig 3A)**. However, this difference disappeared over six months.

**Figure 3.**
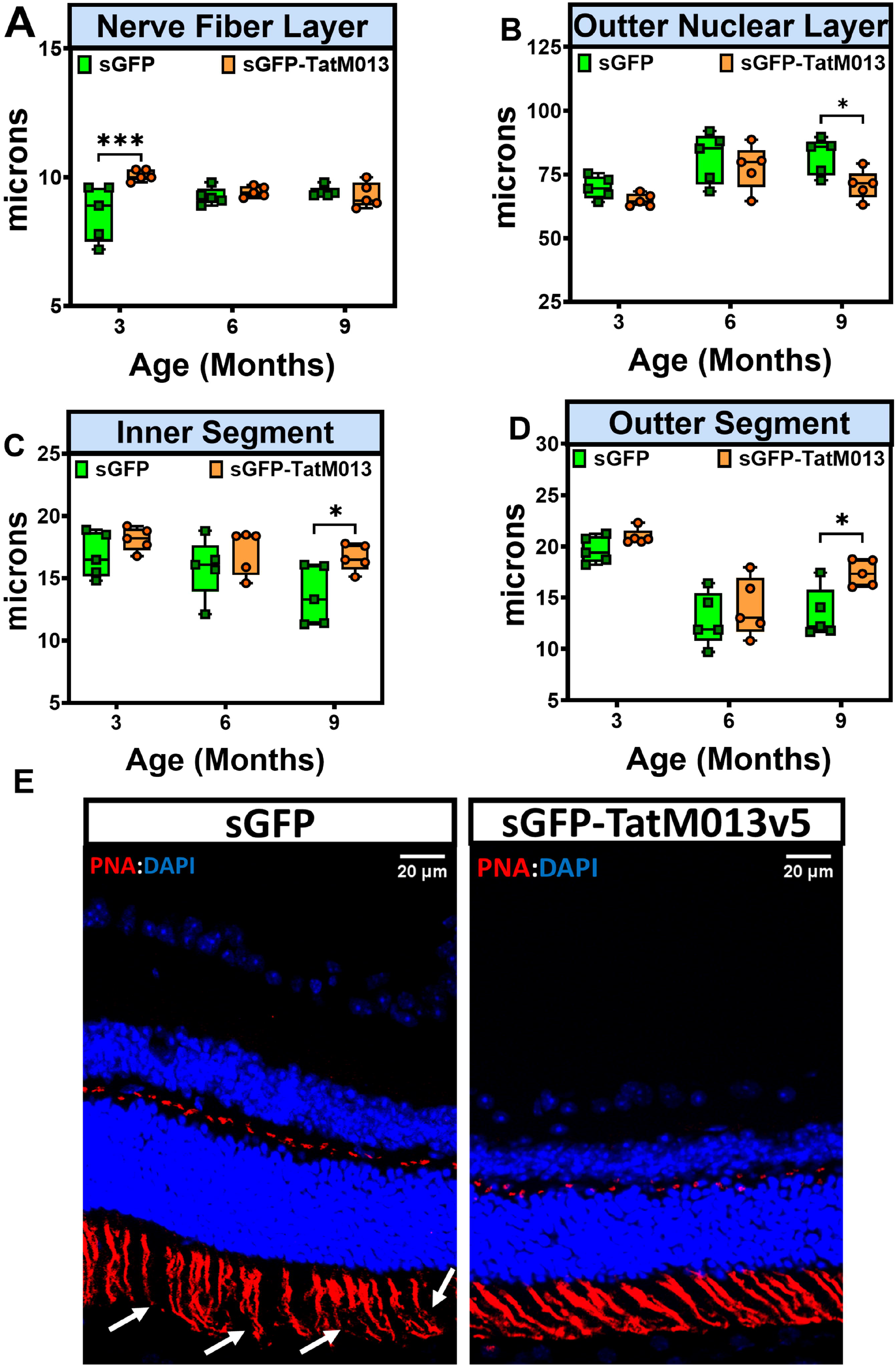
Effects of TatM013 on the retina structure of the RPE-specific Sod2 KO mouse model. Two hundred and fifty B-scans of mice retinas were acquired using a Bioptigen spectral-domain optical coherence tomography. We averaged every 10 B-scans to obtain 25 high-resolution images. Bioptigen Diver auto-segmentation software was used to measure the thickness of each retina layer. The thickness of the **(A)** retinal nerve fiber layer (RNFL), **(B)** outer nuclear layer (ONL), **(C)** inner segments (IS), and **(D)** outer segments (OS) of retinas treated with either sGFP (grey) or sGFP-TatM013v5 (black) were plotted as a function of the animal age. **(E)** Retinas cross sections stained with peanut agglutinin (PNA) to detect the structure of IS and OS. sGFP-TatM013v5 treated retinas had more intact IS and OS based on the PNA staining (red channel). Values were reported as average +/-standard deviation (n=5 mice). * = adj. p-val ≤ 0.05, *** = adj. p-val. ≤ 0.001

Similarly, the thickness of the outer nuclear layer (ONL) was quantified in SD-OCT images to estimate the effects of M013 on photoreceptor survival. Although no significant difference was observed between sGFP and M013-treated eyes at 3- or 6-months of age, the sGFP-treated eyes did show a significantly thicker ONL at nine months of age when compared to M013-treated eyes at the same time point **(Fig 3B)**. The eyes treated with the M013 vector had longer inner and outer segments (IS and OS, respectively) than sGFP-treated eyes **(Fig 3C-D)**. Furthermore, when retina cryosections from both groups were stained with PNA (a lectin that stains cone inner and outer retina glycoproteins^27^), we observed that TatM013v5-treated retinas had more intact inner and outer segments **(Fig 3E, white arrows)**, thus supporting our SD-OCT measurements. These results indicate that TatM013v5 retinal expression can protect photoreceptors’ inner and outer segments, which explains the functional effects described earlier.

### TatM013v5 retinal expression decreases M1-monocytic associated genes

The M013 gene is a potent anti-inflammatory protein that blocks the nuclear translocation of the p65 subunit of the NF-kB transcription factor. In addition, M013 interacts with the ASC-1 protein through a different domain and prevents its incorporation into the NLRP3 inflammasome^7^.

We hypothesized that retinal expression of M013 would decrease the inflammatory aspect of our mouse model of GA. Therefore, frozen sections from animals treated with either sGFP or sGFP-TatM013v5 vector were sectioned and stained with anti-CD45 antibody to detect the presence of immune cells within the retina. In sections of three-month-old mice, there were CD45 positive cells in all the ONL and subretinal space only in sGFP-treated eyes **(Fig 4A)**. Such cells were detected only in the inner retina in TatM013v5 treated eyes **(Fig 4B)**. Notably, CD45-positive cells were detected interacting with the RPE layer only in sGFP-treated eyes.

**Figure 4.**
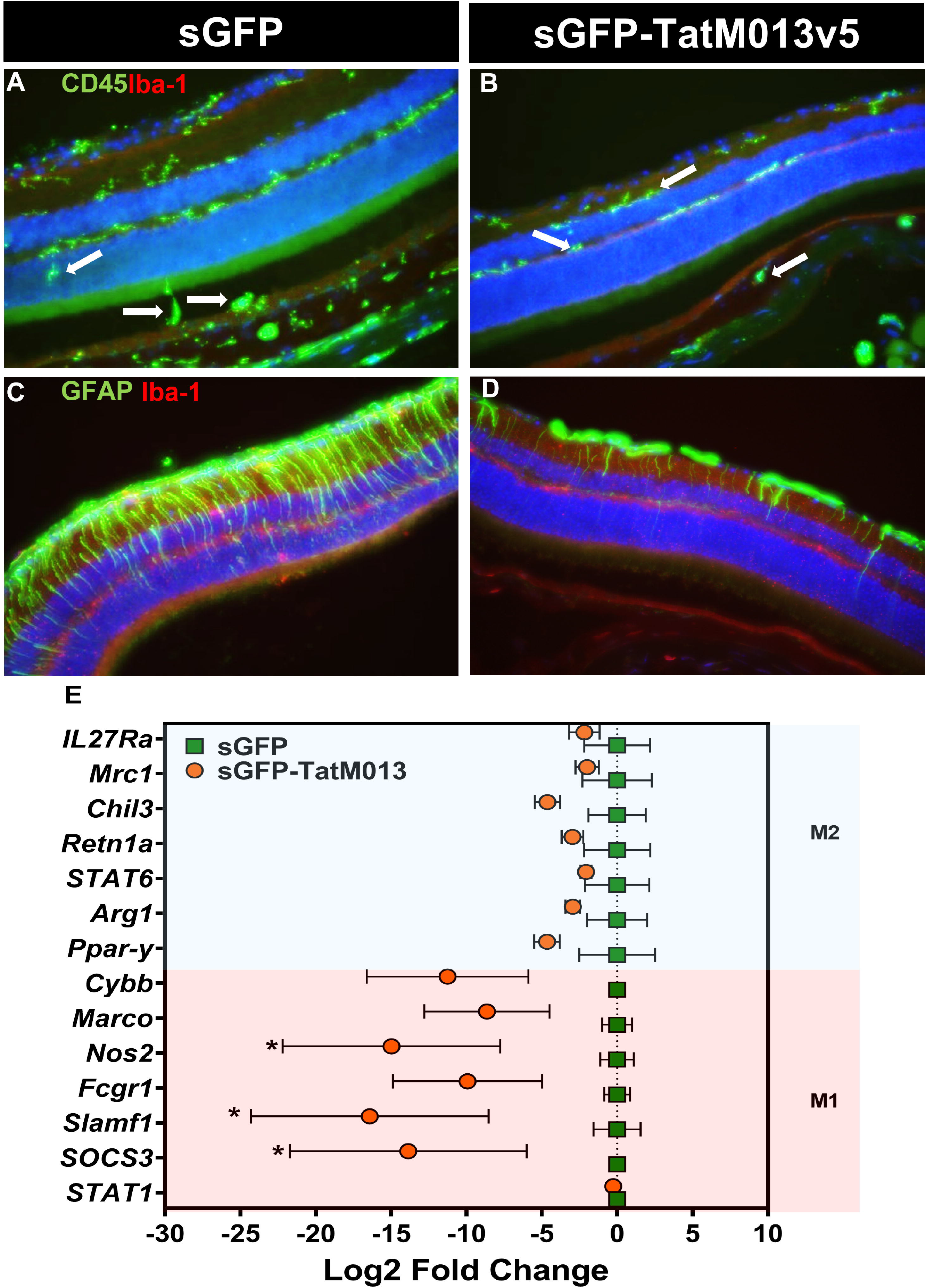
The retinal expression of TatM013v5 decreases reactive gliosis and inflammation. Cryosections from a 3-month-old RPE-specific Sod2 KO mouse model were stained with antibodies against CD45 and Iba-1 or GFAP and Iba-1. **(A)** CD45-positive cells were detected through all the retinal layers in retinas treated with the sGFP vector, especially in the RPE layer (*white arrows*). **(B)** In contrast, retinas treated with the sGFP-TatM013v5 vector showed CD45 positive cells only between the RNFL and the IPL or below the RPE layer (*white arrows*). **(C)** Increased expression of GFAP through all retinal layers was observed in sGFP-treated retinas indicating Muller glial activation. **(D)** The GFAP staining pattern in sGFP-TatM013v5 treated retinas is mainly limited to the RNFL, showing astrocyte expression and minimal staining in Muller glial cells. **(E)** Total RNA was isolated from retinas treated with an AAV vector, delivering either sGFP or sGFP-TatM013v5. Gene expression of M1-associated genes (e.g., STAT1 and Nos2) and M2-associated genes (e.g., PPar-γ and Arg1) were quantified using RT-qPCR. Beta Actin was used as the reference housekeeping gene for all samples. The ΔΔCt method to determine the fold-change. Values are plotted as the Log2 of the fold-change (average ± standard deviation) to illustrate better the decrease in M1-gene expression induced by the expression of sGFP-TatM013v5. No significant changes were identified within the M2 genes. (n=3 retina samples). * = adjusted p-value ≤ 0.05

Next, we studied the activation of Müller glia by immunostaining with an anti-GFAP (glial fibrillary acidic protein). Eyes treated with the sGFP vector showed significant activation of Müller glia based on this staining **(Fig 4C)**. This fluorescent pattern is different from that observed in TatM013v5 treated retinas, which had minor GFAP staining, especially in the transverse orientation, indicative of Müller glia activation **(Fig 4D)**.

Monocytic cells (e.g., microglia, macrophages, and monocytes) are known to express genes involved in either an active inflammatory response (M1 phenotype) or pro-angiogenesis response (M2 phenotype), which are considered anti-inflammatory^28^. To determine how retinal M013 expression impacts the levels of either M1 or M2-associated genes, we used RT-qPCR in retina RNA samples to measure gene expression changes. Expression of M1-related genes was decreased in TatM013v5-treated retinas compared to sGFP-treated retinas **(Fig 4E)**. Specifically, we found a potent inhibition of nitric oxide synthetase 2 (Nos2), signaling lymphocytic activation molecule 1 (Slamf1), and suppressor of cytokine signaling 3 (Socs3). However, no significant effect was observed in M2-related genes, including arginase 1 (Arg1) and peroxisome proliferator-activated receptor gamma (Ppar-γ). These results suggest that retinal expression of TatM013v5 can significantly suppress the pro-inflammatory response associated with our GA mouse model, thus explaining the functional protection of the retina.

### Retinal gene expression of M013 alters retinal cytokine expression

To determine the impact of TatM013v5 retinal expression on the concentration of cytokines and chemokines in the retina, we quantified 32 analytes using multiplex ELISA. Retinal protein extracts were prepared, and equal amounts of proteins were assayed. For simplicity, we compared all the analytes concentrations to their respective concentration at three months **(Fig 5A)**. IL-10, IL-9, and leukemia inhibitory factor (LIF) concentrations were differentially expressed out of the 32 analytes. However, the concentration of these analytes decreased to the same levels as the sGFP control by nine months **(Fig 5B-D)**. Although not statistically significant, we also identified a trend towards an increase in IL-13 in the TatM013v5 treated retinas at three months. Interestingly, the signaling pathways of IL-9, IL-10, LIF, and IL-13 are known to activate STAT3, which has been reported to have retinal neuroprotective effects^29–32^. Together these results suggest that TatM013v5 inhibition of NF-kB activation in the retina induces retinal neuroprotection potentially through STAT3 activation.

**Figure 5.**
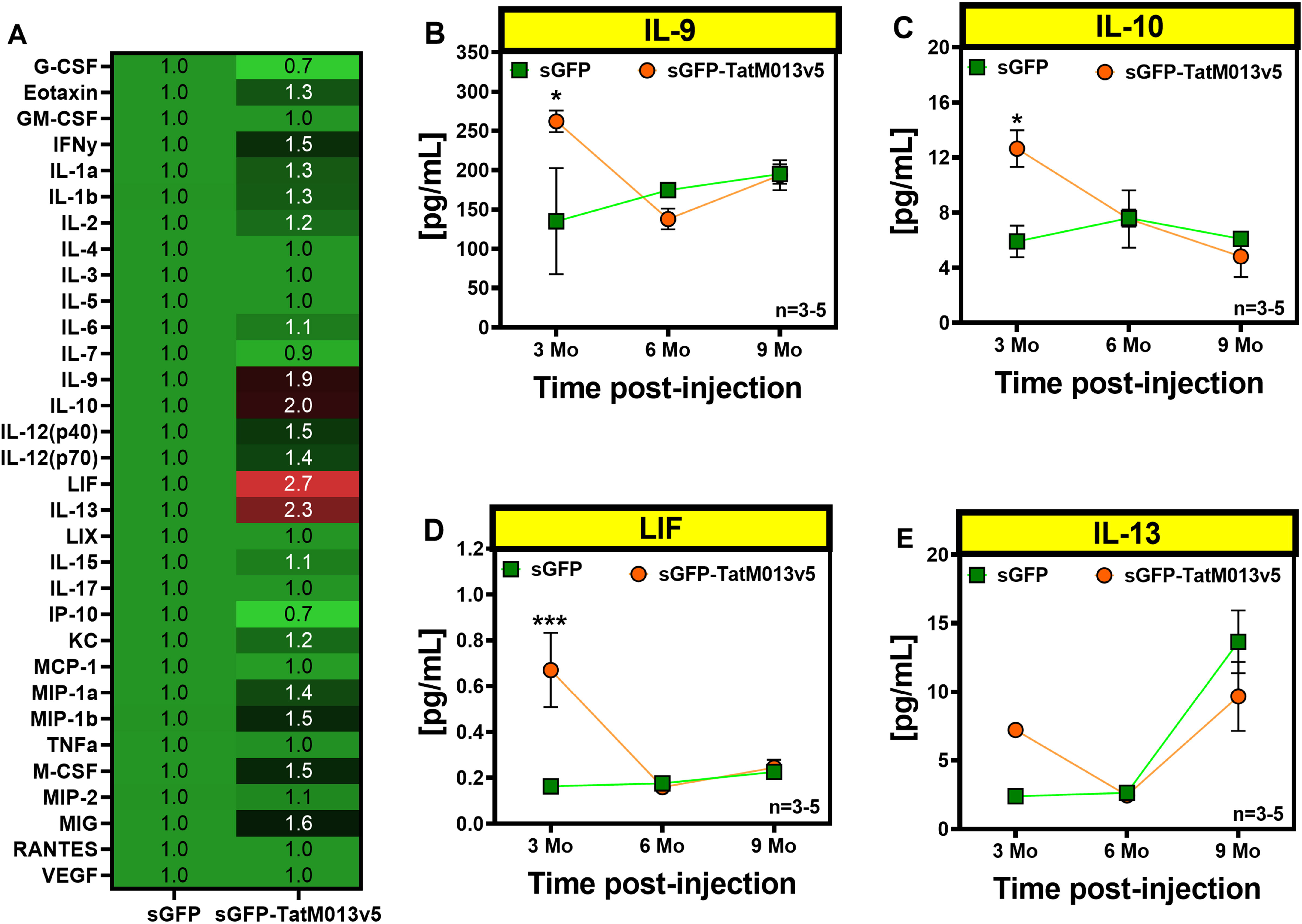
Retinal expression of TatM013v5 increases retinal concentration of IL-10, IL-9, and LIF. Retinal protein extracts were evaluated by multiplex ELISA using a pre-mixed kit for 32 cytokines and chemokines. Four of these analytes were significantly different, at least at a one-time point in the assay. **(A)** At three months of age, the anti-inflammatory cytokine IL-10 is increased in the sGFP-TatM013v5-treated retinas compared to sGFP-treated retinas. **(B)** IL-9 is significantly increased in sGFP-TatM013v5-treated retinas at three months of age. **(C)** Like IL-10, leukemia inhibitory factor (LIF) is increased substantially in sGFP-TatM013v5 treated retinas at three months. **(D)** Finally, the granulocyte colony-stimulating factor (G-CSF) concentration increased in the sGFP-treated retinas by nine months and not in the sGFP-TatM013v5. Values represent average ± standard deviation (n=3-5 retina lysates). * = adj. p-val ≤ 0.05

## DISCUSSION

Our study demonstrates that AAV-delivery of an anti-inflammatory viral gene can curtail the retinal damage induced by chronic RPE oxidative damage in a mouse model of GA. A single intravitreal injection of our AAV vector delivering a secretable and cell-penetrating M013 tagged with a V5 moiety decreased retinal inflammation in the EIU mouse model, similar to the non-tagged secretable and cell-penetrating M013 we have previously described^8^. When injected into the RPE-specific Sod2 cKO mouse model of GA, this vector preserved the amplitudes of the a-, b-, and c-waves from two to nine months of age compared with control. More importantly, the average amplitudes at 9 months among M013-treated mice were similar to or higher than those observed in sGFP-treated animals at 3 months. In addition, TatM013v5 treatment blocked retinal activation of microglia and Müller glia which was observed in control treated eyes. The significant decrease in the expression of M1-associated monocytic genes within these retina samples further validates the anti-inflammatory effects of TatM013v5. These changes in the sGFP-TatM013v5-treated retinas were accompanied by longer photoreceptors inner and outer segments compared to sGFP-treated retinas. Finally, expression of sGFP-TatM1013v5 in the retina of this mouse model increased IL9, IL-10, and LIF compared to sGFP-treated retinas. Together, our results indicate that retinal M013-mediated inhibition of RelA/p65 decreases retinal inflammatory pathways and stimulates neuroprotective pathways through an uncharacterized mechanism.

AMD is a multifactorial disease linked to complex genetics and environmental factors. Variations in the CFH and CFB genes, the presence of monocytic cells in the retina, and complement deposition indicate chronic retinal inflammation and complement-activation are associated with AMD^33,34^. These observations have sparked the interest in developing anti-inflammatory therapies. A clinical study conducted by Jaffe et al. demonstrated that a monthly injection of a C5 inhibitor in AMD patients significantly slowed disease progression using differences in fundus autofluorescence *in vivo*^35^. Unfortunately, this group also found an increase in the incidence of macular neovascularization among patients treated with the C5a inhibitor. Similarly, a clinical trial using a C3 inhibitor in AMD patients showed a significant reduction in the GA area, however it also showed a similar increased incidence in macular neovascularization^36^. These observations support the use of biologicals targeting inflammatory pathways in the retina as potential therapies, but further research is needed to better understand the how inhibition of complement in the retina can lead to neovascularization. Interestingly, we have not observed neovascularization in our mouse model of geographic atrophy. We observed no increase expression of pro-neovascular (M2) genes following sGFP-TatM1013v5 treatment, but we have not determined if prolonged treatment leads to choroidal neovascularization.

The early increase in the expression of LIF, IL-10, and IL-9 suggests a strong activation of STAT3 signaling. Ligand engagement of these cytokines is known to activate STAT3^37–39^. However, how inhibition of RelA activation can lead to increases in these cytokines remains unknown. Gene delivery of a drug tunable form of LIF was shown to significantly slow retinal degeneration in the rd10 mouse model of inherited retinal degeneration^40^. Thus, further understanding of the intricacies of how pro-inflammatory and neuroprotective pathways in the retina interact to maintain homeostasis is needed. Our future studies will address the dynamic mechanism between RelA and STAT3 in the retina as a first step.

## CONCLUSIONS

Retinal inflammation has been recognized as part of the pathobiology of AMD. We have demonstrated that retinal inhibition of p65/RelA and the NLRP3 inflammasome significantly delays loss of retinal function in a mouse model of AMD geographic atrophy form. Our observations support the notion that retinal inflammation is an important disease modifier in dry AMD. Furthermore, inhibiting inflammatory pathways, such as the NF-kB, hold great promise to slow vision loss in the context of dry AMD and indeed can have a great impact in the quality of life of patients afflicted with this disease. Future studies will elucidate the cellular mechanism behind the inhibition of RelA and the delay of retinal degeneration.

## ACKNOWLEDGMENTS

We want to acknowledge the Ocular Gene Therapy Core for generating the AAV vectors used in this work.

## REFERENCES

1. Jonas JB, Cheung CMG, Panda-Jonas S. Updates on the Epidemiology of Age-Related Macular Degeneration. Asia Pac J Ophthalmol (Phila) 2017;6:493–497.

2. Crabb JW. The proteomics of drusen. Cold Spring Harb Perspect Med 2014;4:a017194.

3. Winkler TW, Grassmann F, Brandl C, et al. Genome-wide association meta-analysis for early age-related macular degeneration highlights novel loci and insights for advanced disease. BMC Med Genomics 2020;13:120.

4. Lorés-Motta L, van Beek AE, Willems E, et al. Common haplotypes at the CFH locus and low-frequency variants in CFHR2 and CFHR5 associate with systemic FHR concentrations and age-related macular degeneration. Am J Hum Genet 2021;108:1367–1384.

5. Rahman MM, McFadden G. Myxoma virus lacking the pyrin-like protein M013 is sensed in human myeloid cells by both NLRP3 and multiple Toll-like receptors, which independently activate the inflammasome and NF-κB innate response pathways. J Virol 2011;85:12505–12517.

6. Garg RR, Jackson CB, Rahman MM, et al. Myxoma virus M013 protein antagonizes NF-κB and inflammasome pathways via distinct structural motifs. J Biol Chem 2019;294:8480–8489.

7. Rahman MM, Mohamed MR, Kim M, et al. Co-regulation of NF-kappaB and inflammasome-mediated inflammatory responses by myxoma virus pyrin domain-containing protein M013. PLoS Pathog 2009;5:e1000635.

8. Ildefonso CJ, Jaime H, Rahman MM, et al. Gene delivery of a viral anti-inflammatory protein to combat ocular inflammation. Hum Gene Ther 2015;26:59–68.

9. Ridley RB, Young BM, Lee J, et al. AAV Mediated Delivery of Myxoma Virus M013 Gene Protects the Retina against Autoimmune Uveitis. J Clin Med;8. Epub ahead of print November 29, 2019. DOI: 10.3390/jcm8122082.

10. Marneros AG. Role of inflammasome activation in neovascular age-related macular degeneration. FEBS J. Epub ahead of print November 12, 2021. DOI: 10.1111/febs.16278.

11. Ambati M, Apicella I, Wang S-B, et al. Identification of fluoxetine as a direct NLRP3 inhibitor to treat atrophic macular degeneration. Proc Natl Acad Sci USA;118. Epub ahead of print October 12, 2021. DOI: 10.1073/pnas.2102975118.

12. Allingham MJ, Loksztejn A, Cousins SW, et al. Immunological Aspects of Age-Related Macular Degeneration. Adv Exp Med Biol 2021;1256:143–189.

13. Weaver C, Cyr B, de Rivero Vaccari JC, et al. Inflammasome Proteins as Inflammatory Biomarkers of Age-Related Macular Degeneration. Transl Vis Sci Technol 2020;9:27.

14. Celkova L, Doyle SL, Campbell M. NLRP3 inflammasome and pathobiology in AMD. J Clin Med 2015;4:172–192.

15. Doyle SL, Campbell M, Ozaki E, et al. NLRP3 has a protective role in age-related macular degeneration through the induction of IL-18 by drusen components. Nat Med 2012;18:791–798.

16. Mao H, Seo SJ, Biswal MR, et al. Mitochondrial oxidative stress in the retinal pigment epithelium leads to localized retinal degeneration. Invest Ophthalmol Vis Sci 2014;55:4613–4627.

17. Biswal MR, Han P, Zhu P, et al. Timing of antioxidant gene therapy: implications for treating dry AMD. Invest Ophthalmol Vis Sci 2017;58:1237–1245.

18. Biswal MR, Justis BD, Han P, et al. Daily zeaxanthin supplementation prevents atrophy of the retinal pigment epithelium (RPE) in a mouse model of mitochondrial oxidative stress. PLoS ONE 2018;13:e0203816.

19. Biswal MR, Ildefonso CJ, Mao H, et al. Conditional induction of oxidative stress in RPE: A mouse model of progressive retinal degeneration. Adv Exp Med Biol 2016;854:31–37.

20. Brown EE, DeWeerd AJ, Ildefonso CJ, et al. Mitochondrial oxidative stress in the retinal pigment epithelium (RPE) led to metabolic dysfunction in both the RPE and retinal photoreceptors. Redox Biol 2019;24:101201.

21. Zolotukhin S, Potter M, Zolotukhin I, et al. Production and purification of serotype 1, 2, and 5 recombinant adeno-associated viral vectors. Methods 2002;28:158–167.

22. Ridley RB, Walsh EM, Ildefonso CJ. Molecular design and production of AAV viral vectors for gene therapy. Methods Mol Biol 2021;2225:77–92.

23. Schmittgen TD, Livak KJ. Analyzing real-time PCR data by the comparative C(T) method. Nat Protoc 2008;3:1101–1108.

24. Livak KJ, Schmittgen TD. Analysis of relative gene expression data using real-time quantitative PCR and the 2(-Delta Delta C(T)) Method. Methods 2001;25:402–408.

25. Laemmli UK. Cleavage of structural proteins during the assembly of the head of bacteriophage T4. Nature 1970;227:680–685.

26. Young BM, Jones K, Massengill MT, et al. Expression of a CARD slows the retinal degeneration of a geographic atrophy mouse model. Mol Ther Methods Clin Dev 2019;14:113–125.

27. Long KO, Aguirre GD. The cone matrix sheath in the normal and diseased retina: cytochemical and biochemical studies of peanut agglutinin-binding proteins in cone and rod-cone degeneration. Exp Eye Res 1991;52:699–713.

28. Yadav S, Dwivedi A, Tripathi A. Biology of macrophage fate decision: Implication in inflammatory disorders. Cell Biol Int. Epub ahead of print July 16, 2022. DOI: 10.1002/cbin.11854.

29. Jiang K, Wright KL, Zhu P, et al. STAT3 promotes survival of mutant photoreceptors in inherited photoreceptor degeneration models. Proc Natl Acad Sci USA 2014;111:E5716–23.

30. Rhee KD, Nusinowitz S, Chao K, et al. CNTF-mediated protection of photoreceptors requires initial activation of the cytokine receptor gp130 in Müller glial cells. Proc Natl Acad Sci USA 2013;110:E4520–9.

31. Ueki Y, Wang J, Chollangi S, et al. STAT3 activation in photoreceptors by leukemia inhibitory factor is associated with protection from light damage. J Neurochem 2008;105:784–796.

32. Zhang C, Li H, Liu M-G, et al. STAT3 activation protects retinal ganglion cell layer neurons in response to stress. Exp Eye Res 2008;86:991–997.

33. de Breuk A, Heesterbeek TJ, Bakker B, et al. Evaluating the Occurrence of Rare Variants in the Complement Factor H Gene in Patients With Early-Onset Drusen Maculopathy. JAMA Ophthalmol 2021;139:1218–1226.

34. Heesterbeek TJ, Lechanteur YTE, Lorés-Motta L, et al. Complement activation levels are related to disease stage in AMD. Invest Ophthalmol Vis Sci 2020;61:18.

35. Jaffe GJ, Westby K, Csaky KG, et al. C5 Inhibitor Avacincaptad Pegol for Geographic Atrophy Due to Age-Related Macular Degeneration: A Randomized Pivotal Phase 2/3 Trial. Ophthalmology 2021;128:576–586.

36. Liao DS, Grossi FV, El Mehdi D, et al. Complement C3 Inhibitor Pegcetacoplan for Geographic Atrophy Secondary to Age-Related Macular Degeneration: A Randomized Phase 2 Trial. Ophthalmology 2020;127:186–195.

37. Chucair-Elliott AJ, Elliott MH, Wang J, et al. Leukemia inhibitory factor coordinates the down-regulation of the visual cycle in the retina and retinal-pigmented epithelium. J Biol Chem 2012;287:24092–24102.

38. Donninelli G, Saraf-Sinik I, Mazziotti V, et al. Interleukin-9 regulates macrophage activation in the progressive multiple sclerosis brain. J Neuroinflammation 2020;17:149.

39. Riley JK, Takeda K, Akira S, et al. Interleukin-10 receptor signaling through the JAK-STAT pathway. Requirement for two distinct receptor-derived signals for anti-inflammatory action. J Biol Chem 1999;274:16513–16521.

40. Santiago CP, Keuthan CJ, Boye SL, et al. A Drug-Tunable Gene Therapy for Broad-Spectrum Protection against Retinal Degeneration. Mol Ther 2018;26:2407–2417.

